# Interpreting Rewards from Inverse Reinforcement Learning

**DOI:** 10.64898/2026.07.08.736783

**Authors:** Justin Chow, Yunran Yang, Brokoslaw Laschowski

## Abstract

Inverse reinforcement learning can recover reward functions from observed behavior, but interpreting those rewards remains a fundamental challenge for understanding intelligent behavior and decision-making. To address this challenge, we introduce a novel framework for reward interpretation that combines reward-function analysis, latent mode assignments, and short-history behavioral analysis to infer latent motivations and behavioral dynamics. As a proof-of-concept, we instantiated the framework using switching inverse reinforcement learning on a large-scale dataset of multi-agent social interactions. Our framework interpreted the learned latent modes as ‘cautious’ and ‘volatile’ motivational profiles, demonstrating that recovered reward functions can reveal distinct patterns of behavioral dynamics. More broadly, these findings suggest that the proposed framework provides a promising approach for reverse-engineering and interpreting latent rewards underlying intelligent behavior and decision-making.

## I. Introduction

Understanding behavior and decision-making requires more than predicting actions—it requires inferring the latent rewards that drive them. Reinforcement learning has emerged as a powerful framework for learning from rewards and has been proposed as a general principle underlying intelligent behavior [1]–[5]. Deep reinforcement learning has enabled artificial agents to achieve human-level performance across a variety of tasks [6]–[9]. Inverse reinforcement learning seeks to recover the latent reward functions that best explain behavior [10], [11], providing a computational framework for reverse-engineering internal motivations.

Despite these advances, interpreting reward functions remains a challenge. Inverse reinforcement learning can infer latent rewards from observed behavior, but these are often difficult to interpret, limiting their utility for understanding intelligence. This challenge becomes particularly important in multi-agent social interactions, where behavior is shaped by dynamic interactions between agents and latent motivations may evolve over time. Rather than optimizing for a single objective throughout an interaction, agents may transition between multiple motivational states, each governed by a distinct reward function.

Various inverse reinforcement learning algorithms have been developed to recover latent rewards. For example, maximum entropy inverse reinforcement learning (MaxEnt IRL) by [12] models expert behavior as the maximum-entropy distribution subject to matching observed feature expectations. Although robust to noisy demonstrations, MaxEnt IRL assumes a single, fixed reward function throughout the trajectory, limiting its ability to model changes in motivation over time. Adversarial inverse reinforcement learning (AIRL) developed by [13] introduced an adversarial learning framework capable of recovering transferable reward functions in high-dimensional environments. Although AIRL improves scalability, it also assumes a single underlying reward function throughout each behavioral trajectory.

To address this, Dynamic IRL by [14] introduced continuously varying reward functions by modeling rewards as smooth trajectories over time. This formulation captures gradual changes in latent motivations such as fatigue but is fundamentally Markovian. Consequently, it cannot explicitly represent abrupt transitions between distinct motivational states or account for history-dependent decision-making. Most recently, [15] proposed switching inverse reinforcement learning (SWIRL), which models behavior as transitions between discrete latent modes, each governed by its own reward function and control policy. By introducing both mode switching and history dependency, the algorithm provides a more expressive framework for modeling behavioral sequences. However, its evaluation has focused on structured single-agent environments and interpretation of the reward functions has largely been unexplored.

Here we present a novel framework for interpreting latent reward functions from inverse reinforcement learning (Fig. 1). As a proof of concept, we adapted and applied switching inverse reinforcement learning to a large-scale dataset of multiagent social interactions. Rather than analyzing the recovered rewards in isolation, our framework combines reward-function analysis, latent mode assignments, and short-history behavioral analysis to interpret latent motivations and behavioral dynamics. We demonstrate that our framework can interpret reward functions recovered by inverse reinforcement learning, helping to advance the broader effort of developing more explainable models of intelligence.

**Fig. 1.**
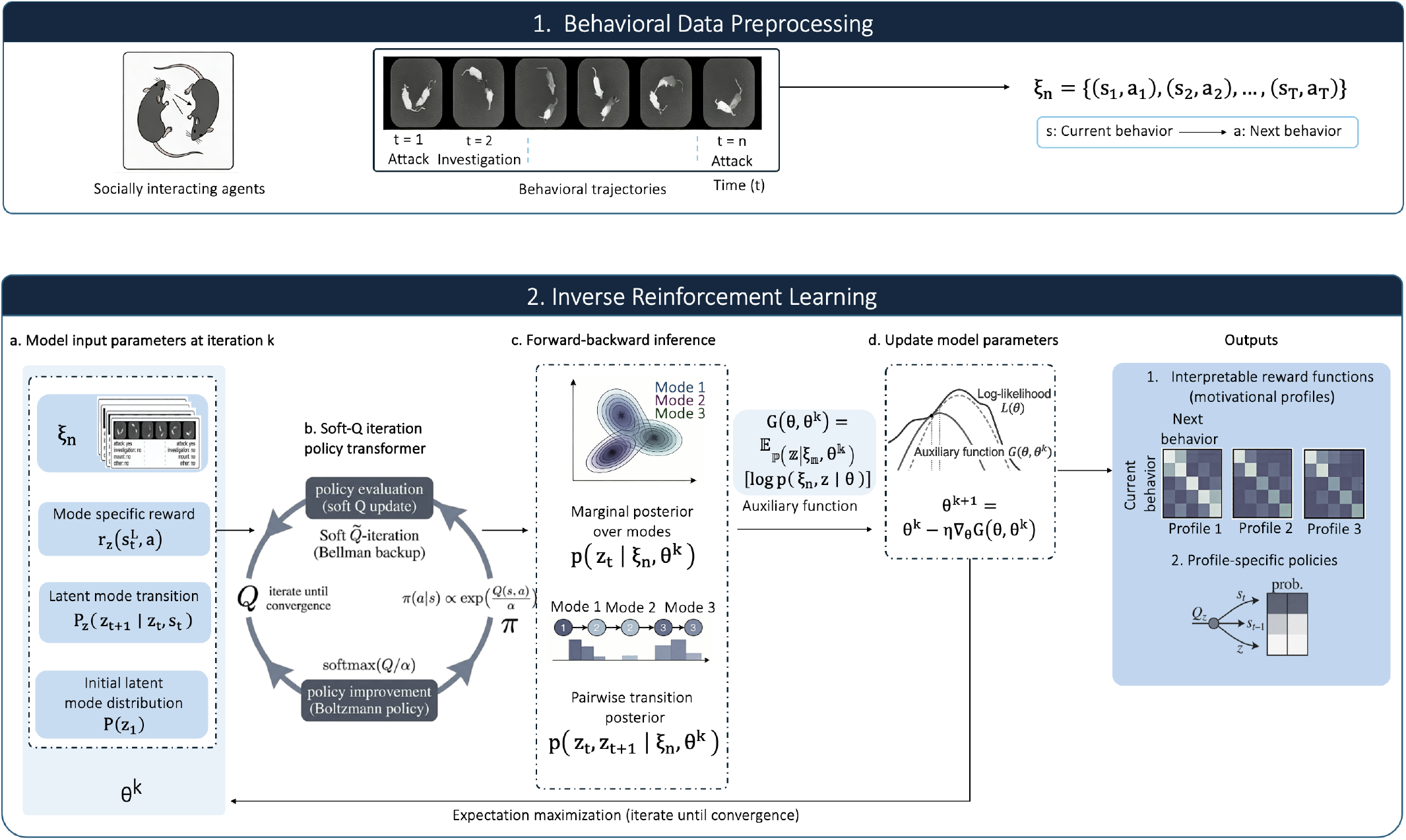
Experimental pipeline used to evaluate our reward interpretation framework. Behavioral trajectories from multi-agent social interactions are preprocessed and analyzed using switching inverse reinforcement learning to recover latent modes, reward functions, and control policies. The recovered reward functions and decoded latent modes are then analyzed through reward-function analysis, latent mode assignments, and short-history behavioral analysis to identify interpretable motivational profiles and characterize behavioral dynamics.

## II. Methods

### A. Inverse Reinforcement Learning

We used switching inverse reinforcement learning (SWIRL) [15] to evaluate our framework. SWIRL models behavior as transitions between latent motivational states, each governed by a distinct reward function and control policy, allowing agents to transition between multiple objectives over time. In this section, we summarize the components of SWIRL that are necessary to understand our reward interpretation framework. At each time step *t*, the agent occupies one of *K* latent motivational states,

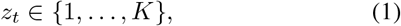

where each latent state is associated with a distinct reward function and control policy. The transition between latent states is conditioned on the current latent state, behavioral state, and action:

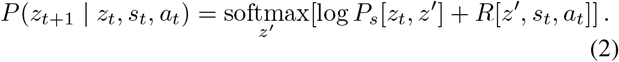

Here, *P*_*s*_ is the base latent-state transition matrix and *R*[*z*^*′*^, *s*_*t*_, *a*_*t*_] is the corresponding reward function. Within each latent state, actions are generated by a mode-specific control policy. To incorporate short-term behavioral history, SWIRL parameterizes the policy as a function of the recent state history 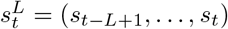:

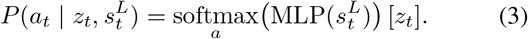

This formulation allows the control policies to depend on recent behavioral context, while the latent modes and reward functions provide mode-specific structure over state-action transitions. Given observed behavioral trajectories, the algorithm jointly estimates the latent mode sequence, reward functions, and control policies by maximizing the likelihood of the observed actions. The joint distribution over latent modes and observed actions is given by

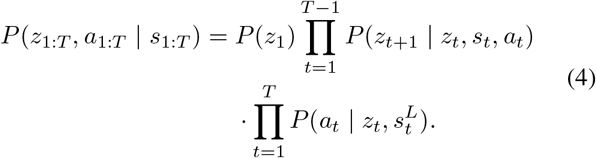

Because the latent mode sequence is unobserved, the algorithm marginalizes over possible mode assignments using posterior mode probabilities. Let *γ*_*t*_ = *P* (*z*_*t*_ | *s*_1:*T*_, *a*_1:*T*_) denote the posterior probability of latent mode *z*_*t*_, and let *ξ*_*t*_ = *P* (*z*_*t*_, *z*_*t*+1_ | *s*_1:*T*_, *a*_1:*T*_) denote the pairwise posterior over consecutive latent modes. The model is trained by maximizing an evidence lower bound:

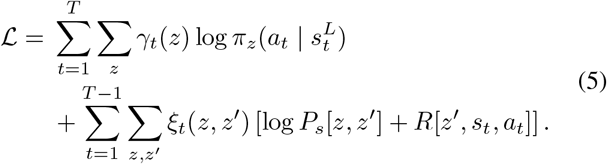

SWIRL is trained using expectation-maximization. During the expectation step, the posterior terms *γ*_*t*_ and *ξ*_*t*_ are estimated using forward-backward message passing. During the maximization step, the model parameters are updated to maximize the evidence lower bound. After convergence, the most likely latent mode sequence is recovered using Viterbi decoding. The recovered reward functions and latent modes form the input to our framework, as subsequently described.

### B. Reward Interpretation Framework

Our reward interpretation framework transforms the latent reward functions from inverse reinforcement learning into interpretable representations of latent motivations and behavioral dynamics. Rather than analyzing the recovered reward functions in isolation, our framework combines reward-function analysis, latent mode assignments, and short-history behavioral analysis to characterize the behavioral preferences associated with each latent motivational state (Fig. 1).

Our framework consists of four steps. First, inverse reinforcement learning is used to recover latent modes, reward functions, and control policies from observed behavior. Second, the reward functions are analyzed as state-action reward matrices to identify the behavioral preferences associated with each latent motivational state. Third, Viterbi-decoded latent mode assignments are examined to characterize how latent motivational states are distributed across behavioral trajectories. Finally, short-history behavioral analysis combines behavioral sequence analysis with the reward functions and control policies to identify interpretable temporal patterns associated with each latent motivational state.

#### 1) Reward Function Analysis

First, the inputs to our framework are the reward functions recovered by inverse reinforcement learning. Each reward function is represented as a state-action reward matrix, where each entry specifies the reward assigned to taking action *a* from behavioral state *s* within a latent motivational state. Rather than interpreting individual rewards in isolation, we analyzed the overall structure of each reward matrix to identify characteristic behavioral preferences. Comparing recovered reward matrices enables the identification of distinct motivational profiles and provides insight into how different reward structures prioritize different behavioral transitions.

#### 2) Latent Mode Assignments

We then analyze the latent mode assignments recovered by SWIRL using Viterbi decoding. The Viterbi algorithm recovers the most likely sequence of latent modes for each behavioral trajectory. Visualization of the decoded latent mode sequences enables qualitative assessment of how latent modes are distributed across behavioral trajectories. We studied whether the latent modes remained stable within individual trajectories or transitioned between modes over time, characterizing the temporal organization of the behavioral dynamics.

#### 3) Short-History Behavioral Analysis

To further characterize the recovered latent modes, we also analyze short-history behavioral sequences associated with each latent mode. While the recovered reward matrices summarize local state-action preferences, they do not directly reveal which recent behavioral histories are most characteristic of each latent mode. Accordingly, we developed a post hoc short-history behavioral analysis that combines behavioral sequence analysis with the recovered reward functions and inferred control policies.

We use an L1-regularized logistic regression model to identify short behavioral sequences whose occurrence in the recent behavioral history is predictive of a target behavior. We selected L1-regularized logistic regression because it provides a sparse and interpretable method for identifying behavioral sequences. Only sequences satisfying a minimum occurrence threshold are retained, and the L1 regularization selects a sparse subset of behavioral sequences for further analysis. For a short behavioral sequence *q* = (*s*_1_, …, *s*_*L*_), latent mode *z*, and target behavior *b*, we compute the target-versus-alternative preference at each sequence position using both the recovered reward function and control policy:

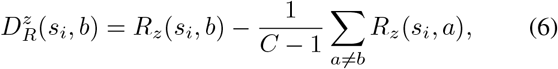

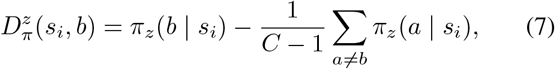

where *C* is the number of behavioral classes. These step-level quantities are averaged across the ordered sequence to obtain reward and policy-derived sequence summaries:

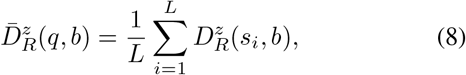

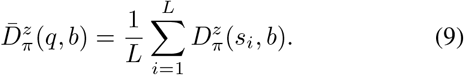

The two sequence summaries are separately normalized using max-absolute scaling and combined to compute a final model-derived ranking value:

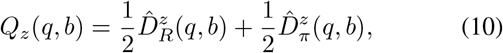

We used equal weighting because there was no prior basis for weighting one source more heavily than the other. Here, 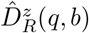 and 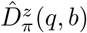 are the normalized reward- and policy-derived sequence summaries, respectively. Higher values of *Q*_*z*_(*q, b*) indicate that the selected behavioral sequence is more strongly associated with the target behavior in latent mode *z*. These ranking values are used to identify characteristic temporal patterns associated with each latent mode and support further interpretation of the motivational profiles.

### C. Experimental Dataset

We evaluated our framework using a large-scale dataset of multi-agent social interactions [16]. The dataset consists of time-series behavioral annotations from top-view videos of freely behaving agents engaged in a resident–intruder assay. In each trial, a resident agent interacted with an intruder agent for 1–10 minutes, with video recorded at 30 Hz. Automated pose estimation was used to track anatomical keypoints, and each frame was manually annotated as one of four behavioral states: *Attack, Investigation, Mount*, or *Other*.

We converted the annotated behavioral states into trajectories for inverse reinforcement learning. Videos with fewer than 2000 frames were discarded, and the remaining videos were truncated to *T* = 2000 frames. This resulted in 58 trajectories, which were partitioned into 46 training trajectories and 12 test trajectories. Each trajectory contained *T*_traj_ = 1999 state-action transitions, where the current behavioral state was treated as *s*_*t*_ and the next behavioral state as *a*_*t*_. Because actions corresponded to transitions to the next behavioral state, the environment was deterministic, with *P* (*s*^*′*^ |*s, a*) = 1 if *s*^*′*^ = *a* and 0 otherwise.

We evaluated both standard and compressed trajectory representations. The standard representation retained all frame-level behavioral transitions, preserving dwell-time information within each behavioral state. The compressed representation removed consecutive self-transitions, retaining only transitions between behavioral states. This reduced each trajectory from 2000 frames to an average of 19 transitions, allowing the model to focus on switching dynamics rather than behavioral persistence.

We also evaluated two state-encoding schemes. In the first-order encoding, each state was represented as a one-hot vector over the four behavioral classes. In the second-order encoding, each state encoded both the current and previous behavioral state using an outer-product representation, yielding a 16-dimensional representation. Across experiments, we varied the number of latent modes *K* = 1, …, 4, the state encoding order, and the trajectory representation to assess their effects on predictive performance and interpretability. Source code for reproducing our experiments and figures are publicly available at: https://github.com/just6660/SWIRL-caltech.

## III. Results

To validate the model, we first measured its ability to predict the next behavioral state across different numbers of latent modes (*K*). Fig. 2 shows the predictive accuracy for the second-order model trained on the compressed dataset, where the model has knowledge of both the current and previous behavioral states when predicting the next action. Training accuracy ranged from 85–88%, while test accuracy ranged from 76–78%, substantially exceeding the 25% chance level expected from randomly selecting among the four behavioral classes. The model with *K* = 2 latent modes achieved the highest test accuracy (78.1%). Increasing the number of latent modes to *K* = 3 or *K* = 4 did not improve generalization, suggesting that two latent modes were sufficient for this particular study. The train-test gap was consistent at roughly 10 percentage points across all values of *K*.

**Fig. 2.**
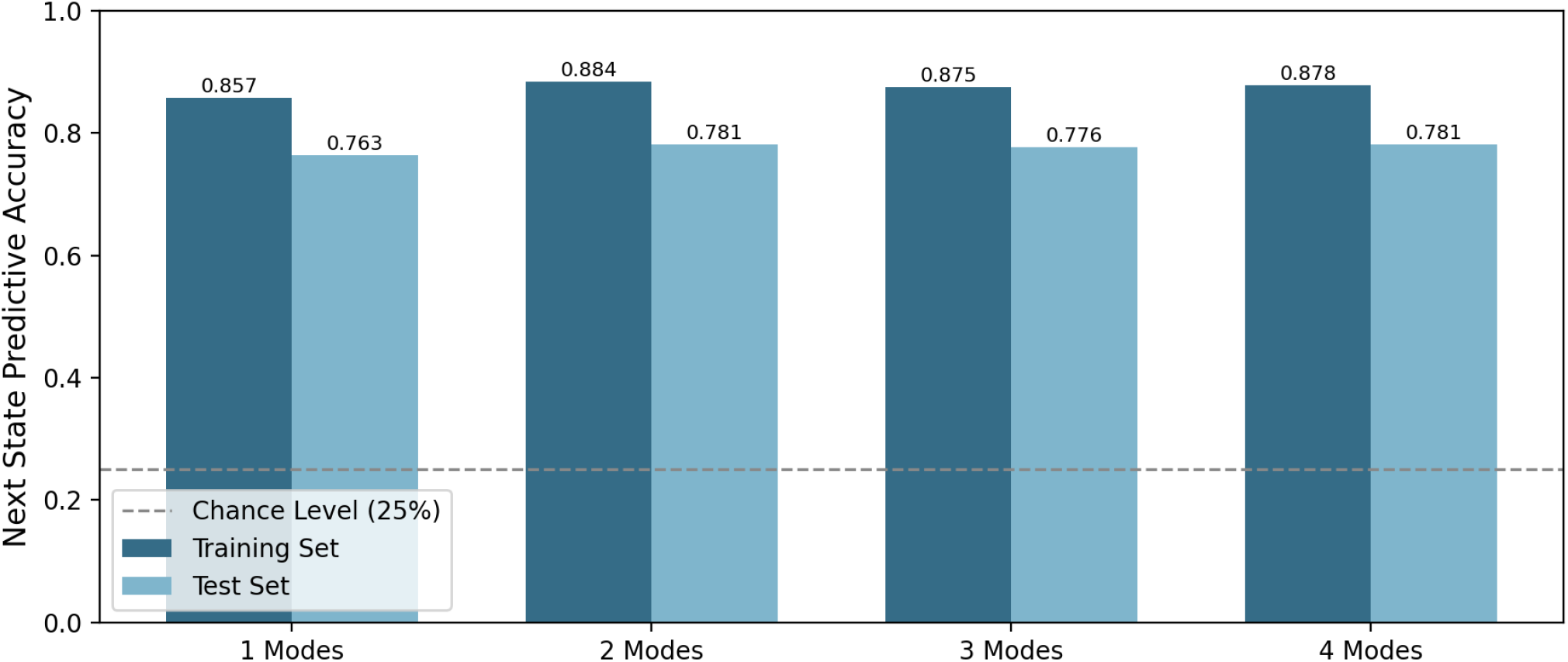
Predictive accuracy on the training and test datasets for the compressed second-order model across different numbers of latent modes (*K* = 1–4). The dashed line denotes the 25% random baseline. The model with *K* = 2 achieved the highest test accuracy.

Next, we analyzed the recovered reward functions from the best-performing model (*K* = 2, compressed dataset). Fig. 3 shows the recovered state-action reward matrices for the two learned latent modes. Each matrix summarizes the reward assigned to transitioning from the current behavioral state (rows) to the next behavioral state (columns). The recovered reward functions exhibited distinct structures, suggesting different underlying motivational strategies.

**Fig. 3.**
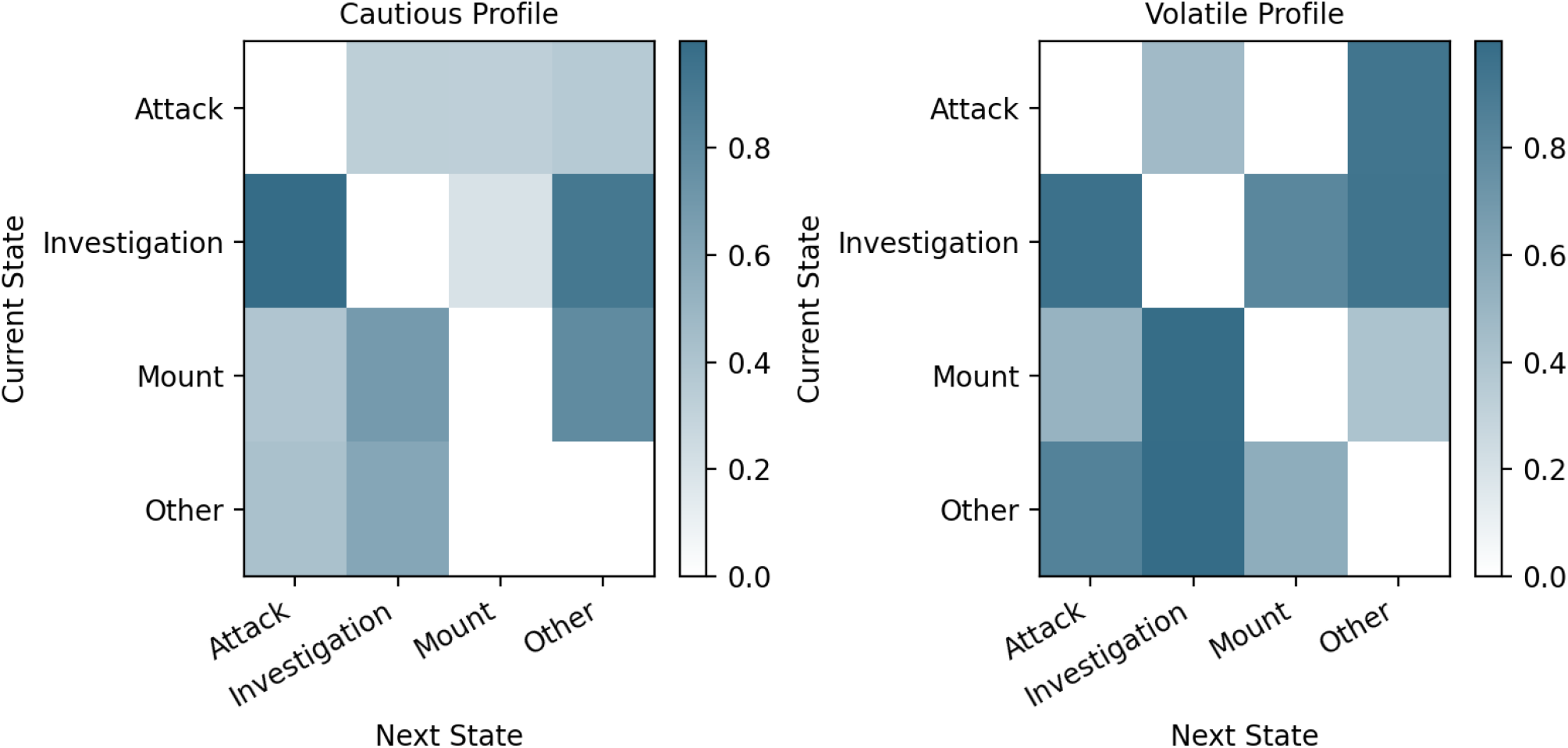
Recovered state-action reward matrices for the two learned latent modes using the *K* = 2 model on the compressed dataset. Rows are the current behavioral state, columns are the next behavioral state, and each entry represents the recovered reward associated with the corresponding state transition. The reward functions exhibit distinct structures, providing the basis for interpreting the latent motivational profiles.

We interpreted the first learned mode as a ‘cautious’ motivational profile in which the reward function consistently assigned low rewards to transitions into *Attack*, especially from the *Investigation* and *Mount* states, while assigning relatively higher rewards to non-aggressive behavioral transitions. In contrast, we interpreted the second learned mode as a ‘volatile’ motivational profile in which the reward function assigned high rewards to transitions into *Attack* from nearly all previous behavioral states, indicating a stronger preference for aggressive behavioral transitions.

We then examined the latent mode assignments by the best-performing model using Viterbi decoding. Fig. 4 shows the decoded latent mode assignments for the *K* = 2 model, where each row represents a behavioral trajectory and the color indicates the inferred latent mode at each transition. The latent mode assignments exhibited strong temporal consistency within individual trajectories. Rather than frequently switching between latent modes, most behavioral trajectories remained assigned to a single latent mode throughout the interaction. Combined with the recovered reward functions, these latent mode assignments suggest that many interactions were reliably characterized by one of the interpreted motivational profiles by our framework.

**Fig. 4.**
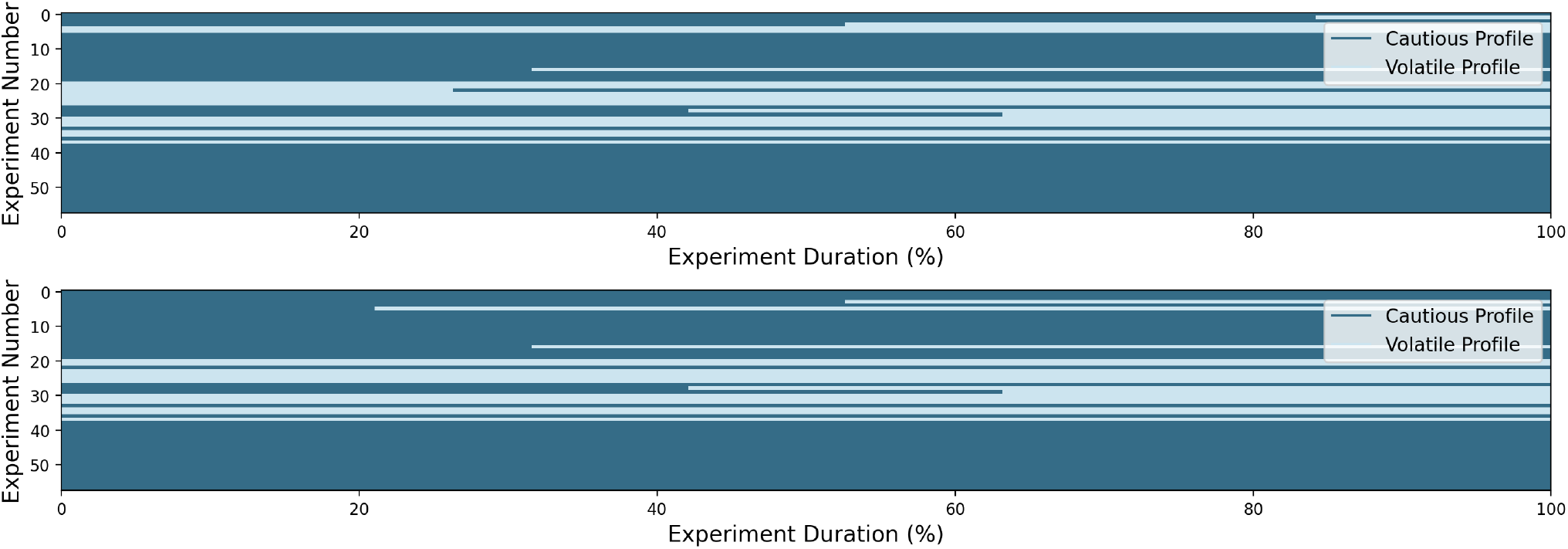
Viterbi-decoded latent mode assignments for the *K* = 2 model on the compressed dataset. Each row represents a behavioral trajectory and the colors denote the inferred latent mode at each transition. Most trajectories remained assigned to a single latent mode throughout the interaction, indicating strong temporal consistency in the latent mode assignments.

Next, we applied the short-history behavioral analysis to identify temporally-extended behavioral patterns associated with each motivational profile. Whereas the recovered reward functions summarized local state-action preferences and the latent mode assignments revealed stable motivational profiles, the short-history analysis provides additional insight into how recent behavioral history influences those preferences. Because transitions into *Attack* most clearly distinguished the two motivational profiles, we focused the short-history behavioral analysis on sequences associated with *Attack*. Fig. 5 shows the ranking values for four categories of short-history behavioral sequences. Higher ranking values indicate a stronger preference for transitions into *Attack*, whereas lower values indicate weaker preference relative to other behaviors.

**Fig. 5.**
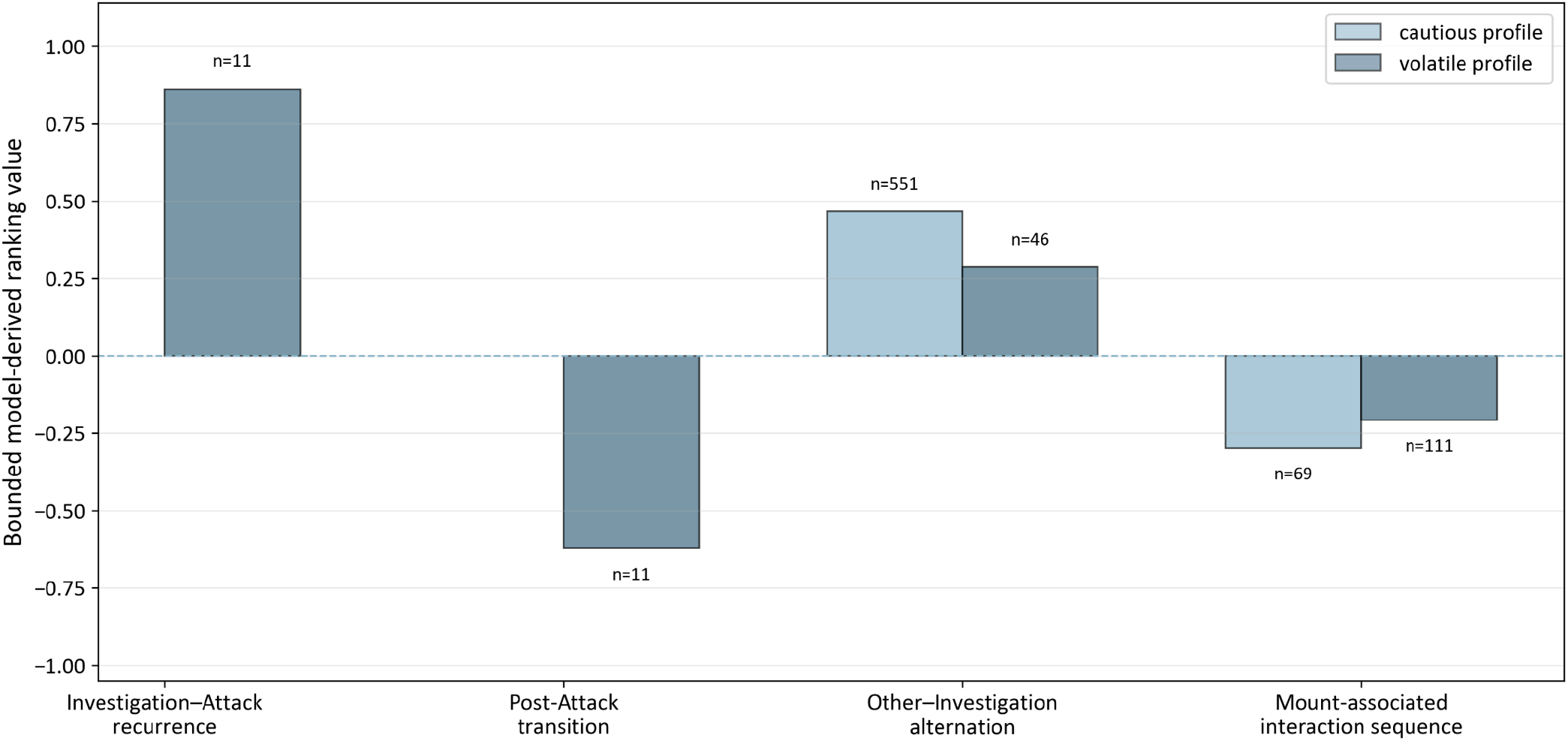
Ranking values for four categories of short-history behavioral sequences. Higher values indicate a stronger preference for transitions into *Attack*. The short-history behavioral analysis identified temporally-extended behavioral patterns that complement the recovered reward functions and further refine the cautious and volatile motivational profiles interpreted by our framework.

The cautious profile showed frequent *Other*–*Investigation* alternation and contained no *Investigation*–*Attack* recurrence, which is consistent with the recovered reward functions indicating less preference for aggressive behavioral transitions. In contrast, the volatile profile showed a strong preference for *Investigation*–*Attack* recurrence, suggesting that repeated escalation within an ongoing interaction was characteristic of this profile. Post-*Attack* transitions showed lower ranking values, suggesting that transitions back to *Investigation* after *Attack* were not characteristic of the volatile profile. *Mount*-associated sequences showed consistently low ranking values across both profiles, suggesting that histories containing *Mount* were generally associated with reduced preference for aggressive behavioral transitions. Together, these findings demonstrate that the short-history behavioral analysis complements the recovered reward functions by identifying temporally-extended behavioral patterns that further refine the motivational profiles interpreted by our framework.

Finally, we examined the effects of trajectory preprocessing. A challenge when applying inverse reinforcement learning to behavioral data is the prevalence of self-transitions. Because the dataset we used was sampled at 30 Hz, the behavioral state at time *t* + 1 was often identical to the behavioral state at time *t*, making self-transitions highly prevalent. When the model was trained on the uncompressed trajectories, it rapidly converged to high log-likelihoods (Fig. 6). However, this mainly reflected learning the persistence of behavioral states rather than the underlying switching dynamics between behaviors. Consistent with this, the Viterbi-decoded latent mode assignments (Fig. 7) were fragmented and exhibited frequent switching between latent modes, preventing us from recovering stable motivational profiles.

**Fig. 6.**
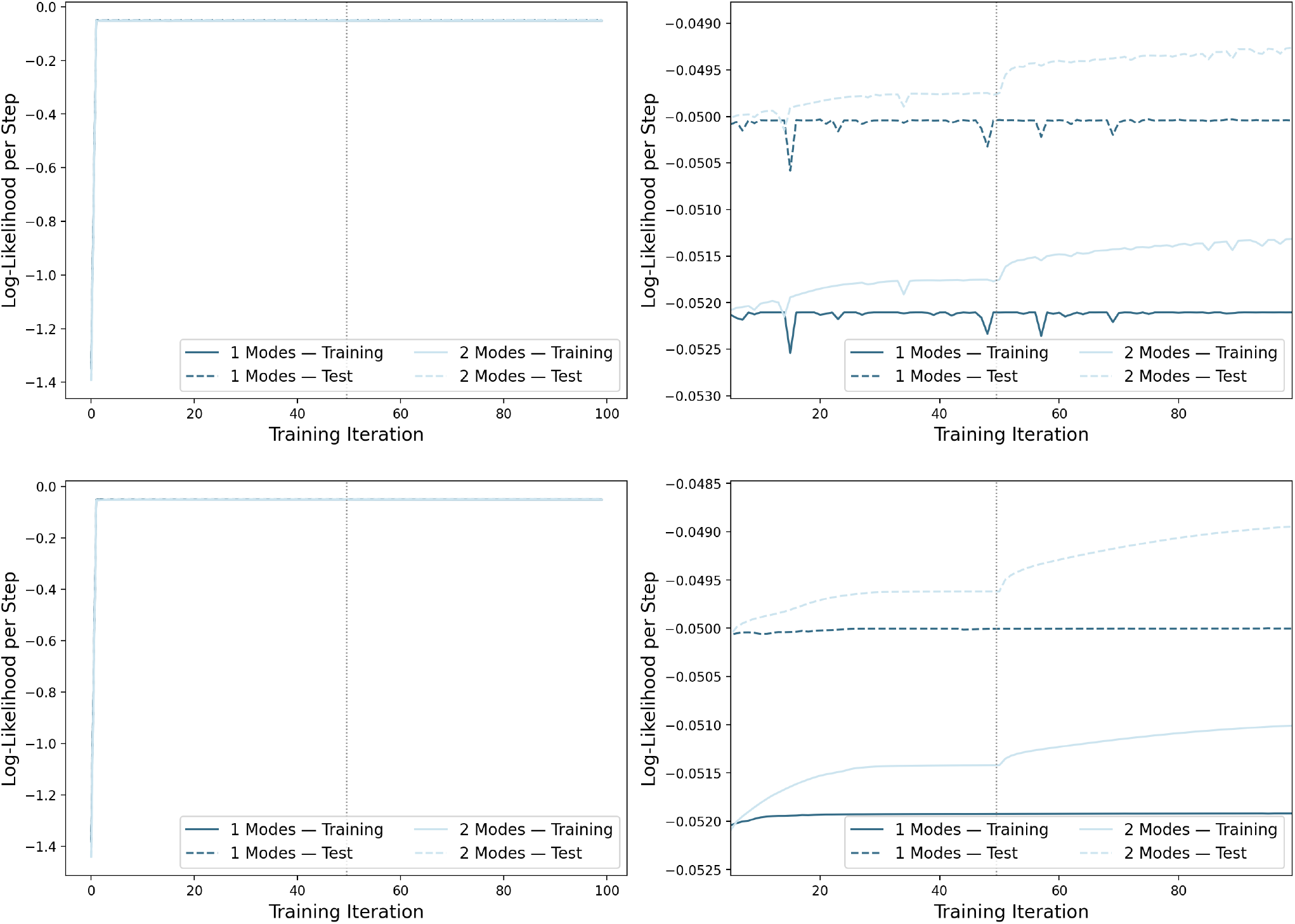
Training and test log-likelihoods for the uncompressed dataset. Both the first-order (top) and second-order (bottom) models achieved high log-likelihoods by exploiting the prevalence of self-transitions rather than learning meaningful switching dynamics.

**Fig. 7.**
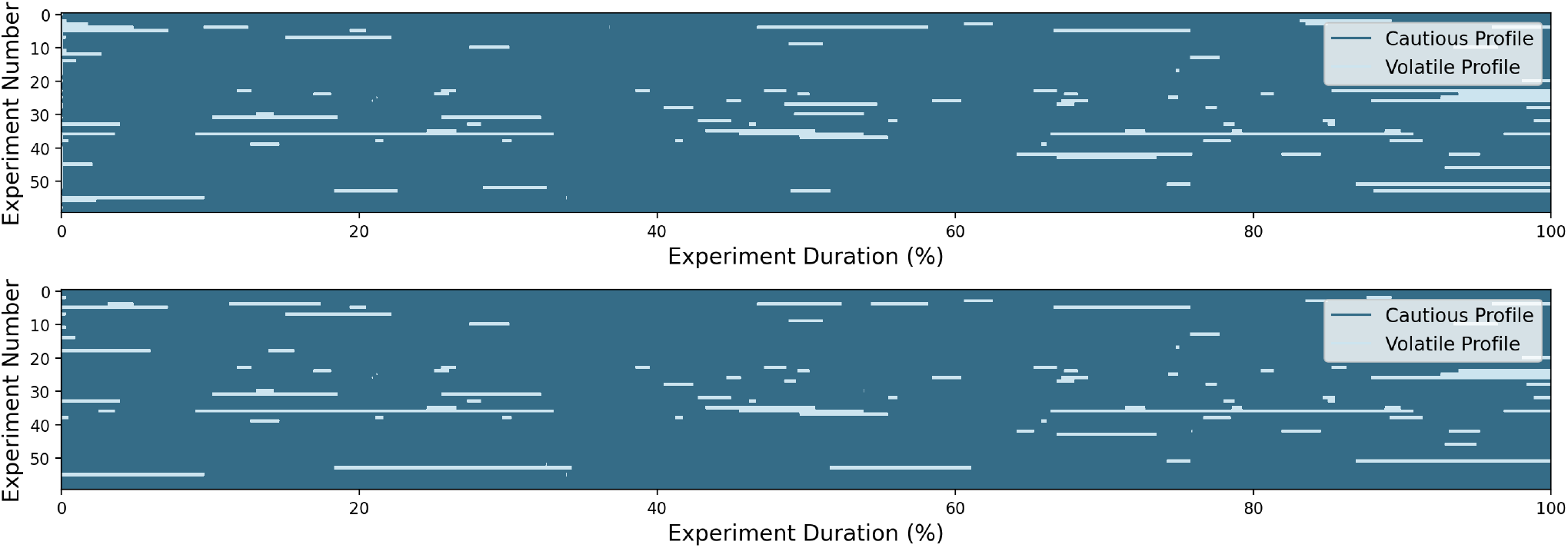
Viterbi-decoded latent mode assignments for the *K* = 2 model on the uncompressed dataset. Each row represents a behavioral trajectory and the colors denote the inferred latent mode at each transition. The latent mode assignments were fragmented and exhibited frequent switching between latent modes, indicating weak temporal consistency compared with the compressed dataset (Fig. 4).

In contrast, removing consecutive self-transitions forced the model to focus on transitions between behavioral states rather than behavioral persistence. This preprocessing produced substantially more stable latent mode assignments (Fig. 4), enabling recovery of interpretable reward functions and motivational profiles. These findings demonstrate that trajectory preprocessing is an important consideration when applying inverse reinforcement learning to natural behavioral data.

## IV. Discussion

In this study, we introduced a novel framework for interpreting reward functions recovered by inverse reinforcement learning. As a proof-of-concept, we adapted and applied switching inverse reinforcement learning to a large-scale dataset of multi-agent social interactions. Our framework combined reward-function analysis, latent mode assignments, and short-history behavioral analysis to characterize latent motivations and behavioral dynamics. Using this framework, we interpreted the learned latent modes as ‘cautious’ and ‘volatile’ motivational profiles, demonstrating that reward functions can reveal distinct patterns in behavioral dynamics. More broadly, these findings demonstrate that our framework offers a promising method for reverse-engineering and interpreting reward functions underlying intelligent behavior and decision-making, complementing other computational approaches that explain behavior using optimization and control theory [17].

Our framework extends prior inverse reinforcement learning research by shifting the focus from reward recovery to reward interpretation. For example, MaxEnt IRL provides a probabilistic framework for recovering a single reward function from observed behavior under a maximum-entropy trajectory distribution [12]. Dynamic IRL extends this by allowing rewards to vary continuously over time, parameterizing the reward as a time-varying combination of spatial reward maps [14]. SWIRL further extends this line of research by modeling behavior as transitions between discrete latent modes with history-dependent reward functions and control policies [15]. Our contribution, in contrast, is not a new inverse reinforcement learning algorithm, but rather a framework for interpreting reward functions recovered by such algorithms. By integrating multiple complementary analyses, our framework systematically interprets the recovered reward functions, identifies motivational profiles, and relates them back to observable behavioral patterns.

Despite these contributions, several limitations remain. First, our framework was evaluated using switching inverse reinforcement learning, which represents behavioral history using finite-order state encoding, limiting its ability to capture long-range temporal dependencies. Consistent with this limitation, our second-order model achieved higher training performance without improving generalization, suggesting increased model complexity without corresponding gains in predictive performance. Second, although our framework improves interpretability, the underlying reward functions and control policies remain difficult to understand mechanistically. The proposed short-history behavioral analysis provides additional insight into the learned model, but it does not explain how behavioral history is internally represented or how those learned representations give rise to the reward function. Finally, our evaluation was performed on a single dataset using a coarse set of human-annotated behavioral classes. Although this provides a useful proof-of-concept, further evaluation across more diverse behavioral domains is necessary to establish the generality of our framework.

These limitations motivate directions for future research. Incorporating Transformer-based encoders into inverse reinforcement learning may enable modeling longer behavioral histories without increasing the state representation [18], [19]. Combining these model architectures with interpretable analyses of reward functions may further improve understanding of how behavioral history shapes decision-making. Moreover, integrating our reward interpretation framework with emerging approaches in neural representational alignment [20], [21] may provide new opportunities to compare representations learned by inverse reinforcement learning with those underlying biological intelligence. Similar opportunities may arise from studies connecting reinforcement learning with computational neuroscience [22]–[24].

## Acknowledgments

We dedicate this research to the students and researchers in Ukraine. Their resilience and unwavering commitment to education and learning continue to serve as a beacon of hope and inspiration to the global academic community.

